# Algorithmic differentiation improves the computational efficiency of OpenSim-based optimal control simulations of movement

**DOI:** 10.1101/644245

**Authors:** Antoine Falisse, Gil Serrancolí, Christopher L. Dembia, Joris Gillis, Friedl De Groote

## Abstract

Algorithmic differentiation (AD) is an alternative to finite differences (FD) for evaluating function derivatives. The primarily aim of this study was to demonstrate the computational benefits of using AD instead of FD in OpenSim-based optimal control simulations. The secondary aim was to evaluate computational choices including different AD tools, different linear solvers, and the use of first- or second-order derivatives. First, we enabled the use of AD in OpenSim through a custom source code transformation tool and through the operator overloading tool ADOL-C. Second, we developed an interface between OpenSim and CasADi to perform optimal control simulations. Third, we evaluated computational choices through simulations of perturbed balance, two-dimensional predictive simulations of walking, and three-dimensional tracking simulations of walking. We performed all simulations using direct collocation and implicit differential equations. Using AD through our custom tool was between 1.8 ± 0.1 and 17.8 ± 4.9 times faster than using FD, and between 3.6 ± 0.3 and 12.3 ± 1.3 times faster than using AD through ADOL-C. The linear solver efficiency was problem-dependent and no solver was consistently more efficient. Using second-order derivatives was more efficient for balance simulations but less efficient for walking simulations. The walking simulations were physiologically realistic. These results highlight how the use of AD drastically decreases computational time of optimal control simulations as compared to more common FD. Overall, combining AD with direct collocation and implicit differential equations decreases the computational burden of optimal control simulations, which will facilitate their use for biomechanical applications.

## Introduction

Combining musculoskeletal modeling and dynamic simulation is a powerful approach to study the mechanisms underlying human movements. There exist two types of model-based dynamic simulations. Inverse simulations identify controls (e.g., muscle excitations) underlying a given movement whereas forward simulations generate the movement based on the controls. Forward simulations are commonly combined with numerical optimization techniques to compute optimal controls. In the resultant optimal control problems, an objective function is minimized subject to constraints describing the musculoskeletal dynamics. Researchers have used such optimal control simulations for tracking measured movements [1–3] or for predicting *de novo* movements [4–7]. Forward simulations have the potential to reveal cause-effect relationships that cannot be explored through inverse simulations. Nevertheless, the nonlinearity and stiffness of the equations describing the dynamic constraints cause the underlying optimal control problems to be challenging to solve and computationally expensive [4,6,7]. For example, small changes in controls can cause large changes in kinematics and hence a foot to penetrate into the ground, drastically increasing ground reaction forces. These challenges have caused the biomechanics community to primarily use simpler inverse simulations.

Over the last decade, the increase in computer performance and the use of efficient numerical methods have equipped researchers with better tools for generating computationally efficient optimal control simulations. In particular, direct collocation methods [3,5,7–10] and implicit formulations of the musculoskeletal dynamics [9,11] have become popular. Direct collocation reduces the sensitivity of the objective function to the optimization variables, compared to other methods such as direct shooting [4], by reducing the time horizon of the integration. Direct collocation converts optimal control problems into large sparse nonlinear programming problems (NLP) that readily available NLP solvers (e.g., IPOPT [12]) can solve efficiently. Implicit formulations of the musculoskeletal dynamics improve the numerical conditioning of the NLP over explicit formulations by, for example, removing the need to divide by small muscle activations [9] or invert a mass matrix that is near-singular due to body segments with a large range of masses and moments of inertia [11]. In implicit formulations, additional controls are typically introduced for the time derivative of the states, which allows imposing the nonlinear dynamic equations as algebraic constraints in their implicit rather than explicit form (i.e., *ẏ* = *u*, 0 = *f_i_*(*y,u*) instead of *ẏ* = *f_e_*(*y*)).

Algorithmic differentiation (AD) is another numerical tool that can improve the efficiency of optimal control simulations [9]. Yet it has been underused in biomechanics. AD is a technique for evaluating derivatives of functions represented by computer programs [13]. It is, therefore, an alternative to finite differences (FD) for evaluating the derivative matrices required by the NLP solver, namely the objective function gradient, the constraint Jacobian, and the Hessian of the Lagrangian (henceforth referred to as simply Hessian). These evaluations are obtained free of truncation errors, in contrast with FD, and for a computational cost of the same order of magnitude as the cost of evaluating the original function. AD relies on the observation that any function can be broken down into a sequence of elementary operations, forming an expression graph (example in Fig 1). AD then relies on the chain rule of calculus that describes how to calculate the derivative of a composition of functions [13]. By traversing a function’s expression graph while applying the chain rule, AD allows computing the function derivatives.

**Fig 1.**
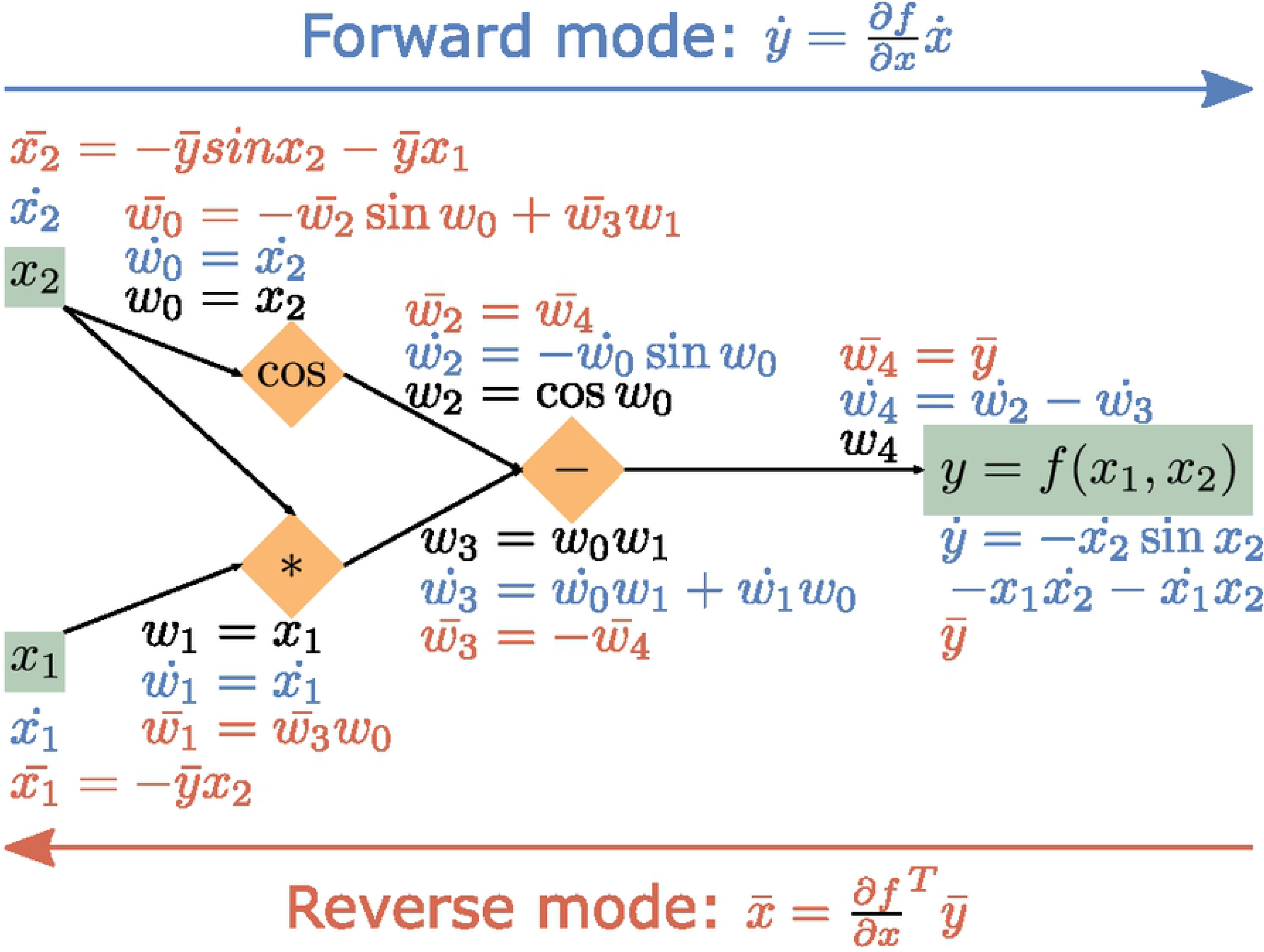
Example of AD forward and reverse modes. A function *y* = *f*(*x*_1_,*x*_2_) = *cos x*_2_ − *x*_1_*x*_2_ is broken down into a sequence of elementary operations, forming an expression graph. In the forward mode, the forward seeds *ẋ*_1_ and *ẋ*_2_ are propagated from the inputs to the output and the Jacobian *J* = *∂f*/*∂**x*** relates *ẋ*_1_ and *ẋ*_2_ and forward sensitivity *ẏ*. In the reverse mode, the reverse seed *ȳ* is propagated from the output to the inputs and the transposed Jacobian *J^T^* relates *ȳ* and reverse sensitivities 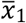 and 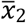.

AD allows traversing the expression graph in two directions or modes: from the inputs to the outputs in its forward mode, and from the outputs to the inputs in its reverse mode. This permits the evaluation of two types of directional derivatives: Jacobian-times-vector product and Jacobian-transposed-times-vector product in the forward and reverse mode, respectively. The computational efficiency of the AD modes depends on the problem dimensions. Consider the function 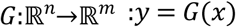 describing the *m* NLP constraints *y* as a function of the *n* optimization variables *x*. The constraint Jacobian *J* = *∂y*/*∂x* is a matrix with size *m x n*. In the forward mode, *J* relates forward seeds *ẋ* and forward sensitivities *ẏ*: *ẏ* = *Jẋ* (example in Fig 1). In the reverse mode, *J^T^* relates reverse seeds *ȳ* and reverse sensitivities 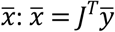 (example in Fig 1). In the forward mode, the cost of evaluating *J* is proportional to *n* times the cost of evaluating *G*. In the reverse mode, the cost of evaluating *J^T^* is proportional to *m* times the cost of evaluating *G*. If there are many more inputs *n* than outputs *m*, the reverse mode may drastically decrease the number of function evaluations required to evaluate *J* and highly reduce the computational (CPU) time as compared to the forward mode [13,14].

OpenSim [15,16] and its dynamics engine Simbody [17] are widely used open-source software packages for musculoskeletal modeling and dynamic simulation but do not leverage tools for AD. Two main approaches exist for adding AD to existing software. In the operator overloading (OO) approach, the AD algorithms are applied after the evaluation of the original function using concrete numerical inputs. This is typically performed by introducing a new numerical type that stores information about partial derivatives as calculations proceed (e.g., through OO in C++) [13,14]. In the source code transformation (SCT) approach, the AD tool analyzes a given function’s source code and outputs a new function that computes the forward or reverse mode of that function. SCT is inherently faster than OO, but may not be readily available for all features of a programming language.

Once a computer program supports AD, the program can be differentiated using the AD algorithms to evaluate the derivative matrices required by the NLP solver. Note that, like FD, AD algorithms can also exploit the sparsity of these matrices resulting from using direct collocation [18]. Of the many open-source AD frameworks, CasADi [14,19] is a modern actively developed tool with many additional features (e.g., code generation) and also interfaces with NLP solvers designed to handle large and sparse NLPs (e.g., IPOPT). CasADi provides a high-level, symbolic, way to construct an expression graph, on which SCT is applied. The resultant expression graph can be code-generated to achieve the computational efficiency of pure SCT.

The contribution of this study is threefold. First, we enabled the use of AD in OpenSim and Simbody (henceforth referred to as OpenSim). We compared two approaches: we incorporated the OO AD tool ADOL-C [20] and we developed our own AD tool Recorder that uses OO to construct an expression graph on which SCT is applied. Second, we interfaced OpenSim with CasADi, enabling optimal control simulations using OpenSim’s multi-body dynamics models while benefitting from CasADi’s efficient interface with NLP solvers. Third, we evaluated the efficiency of different computational choices based on optimal control simulations of varying complexity solved with IPOPT. We compared three different derivative scenarios: AD with ADOL-C, AD with Recorder, and FD. In addition, we compared different linear solvers and different Hessian calculation schemes within IPOPT, to aid users in choosing the most efficient solver settings. Primal-dual interior point methods such as IPOPT rely on linear solvers to solve the primal-dual system, which involves the Hessian, when computing the Newton step direction during the optimization [21]. The Hessian can be exact (i.e., based on second-order derivative information) or approximated with a limited-memory quasi-Newton method (L-BFGS) that only requires first-order derivative information. We found that using AD through Recorder was more efficient than using FD or AD through ADOL-C whereas the efficiency of the linear solver and Hessian calculation scheme was problem-dependent.

## Materials and methods

### Tools to enable the use of AD in OpenSim

We first incorporated the OO AD tool ADOL-C in OpenSim. ADOL-C relies on the concept of active variables, which are variables that may be considered as differentiable quantities at some time during the execution of a computer program [20]. To distinguish these variables and store information about their partial derivatives, ADOL-C introduced the augmented scalar type *adouble* whose real part is of standard type *double*. All active variables should be of type *adouble*. To differentiate OpenSim functions using ADOL-C, we modified OpenSim’s source code by replacing the type of potential active variables to *adouble* (example for SimTK::square() in Fig 2). We maintained a layer of indirection so that OpenSim could be compiled to use either *double* or *adouble* as the scalar type. We excluded parts of the code, such as numerical optimizers, that were not relevant to this study.

**Fig 2.**
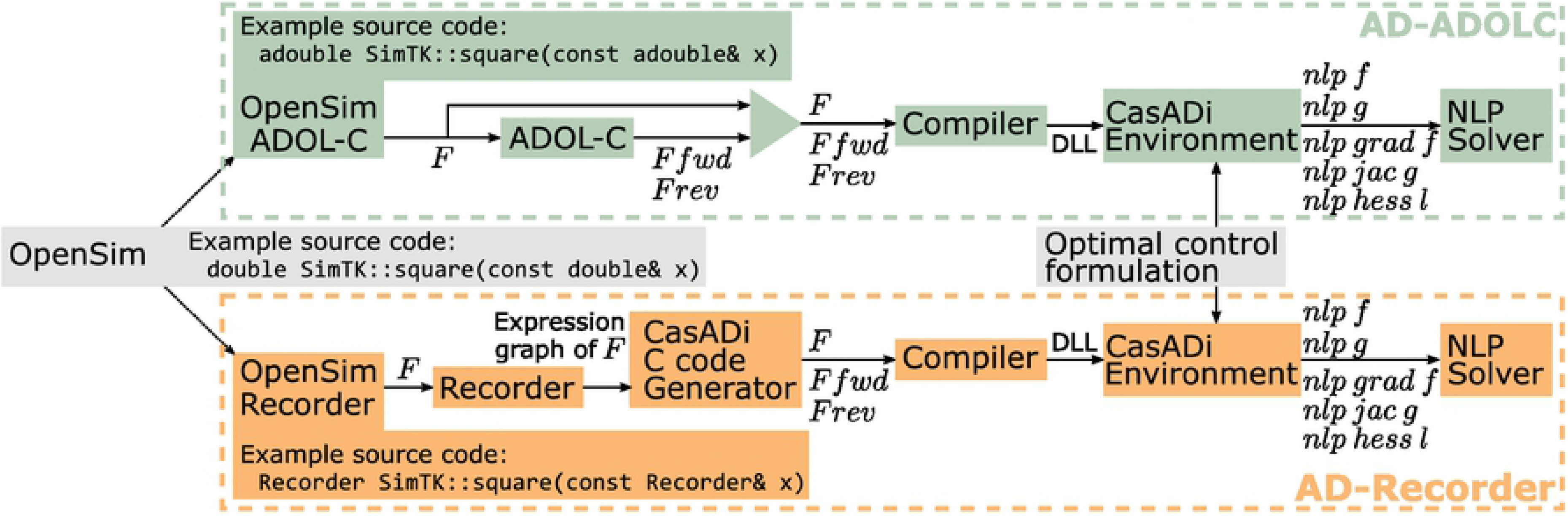
Flowchart depicting the optimal control framework. We developed two approaches (AD-ADOLC and AD-Recorder) to make an OpenSim function *F* and its forward (*F fwd*) and reverse (*F rev*) directional derivatives available within the CasADi environment for use by the NLP solver during the optimization. In the AD-ADOLC approach (top), ADOL-C’s algorithms are used in a C++ code to provide *F fwd* and *F rev*. In the AD-Recorder approach (bottom), Recorder provides the expression graph of *F* as MATLAB source code from which CasADi’s C-code generator generates C-code containing *F*, *F fwd*, and *F rev*. The AD-Recorder approach combines OO, when generating the expression graph, and SCT, when processing the expression graph to generate C-code for *F*, *F fwd*, and *F rev*. In both approaches, the code comprising *F*, *F fwd*, and *F rev* is compiled as a Dynamic-link Library (DLL), which is imported as an external function within the CasADi environment. In our application, *F* represents the multi-body dynamics and is called when formulating the optimal control problem. The latter is then composed into a differentiable optimal control transcription using CasADi. During the optimization, CasADi provides the NLP solver with evaluations of the NLP objective function (*nlp f*), constraints (*nlp g*), objective function gradient (*nlp grad f*), constraint Jacobian (*nlp jac g*), and Hessian of the Lagrangian (*nlp hess l*). CasADi efficiently queries *F fwd* and *F rev* to construct the full derivative matrices.

The limited computational benefits of using AD through ADOL-C led us to seek alternative AD strategies (see discussion for more detail). We developed our own tool, Recorder, which combines the versatility of OO and the speed of SCT. Recorder is a C++ scalar type for which all operators are overloaded to generate an expression graph. When evaluating an OpenSim function numerically at a nominal point, Recorder generates the function’s expression graph as MATLAB source code in a format that CasADi’s AD algorithms can transform into C-code (see S1 Appendix for source code of example from Fig 1). Note that this workflow is currently only practical when the branches (*if-tests*) encountered at the nominal point remain valid for all evaluations encountered during the optimization.

To use Recorder with OpenSim, we relied on the code we had modified for incorporating ADOL-C but replaced *adouble* with the *Recorder* scalar type (example for SimTK::square() in Fig 2). This change required minimal effort but enabled Recorder to identify all differentiable variables when constructing the expression graphs.

### Interface between OpenSim and CasADi

We made utilizable an OpenSim function within the CasADi environment by compiling the function and its derivatives as a Dynamic-link Library (DLL) that then is imported as an external function for use by CasADi (Fig 2). The function derivatives can be computed through ADOL-C (AD-ADOLC in Fig 2) or through Recorder (AD-Recorder in Fig 2).

### Optimal control simulations to evaluate computational choices

We designed three example optimal control simulations to evaluate different computational choices (see Table 1 for detailed formulations). The simulations are optimal control problems whose general formulation consists of computing the controls ***u***(*t*), states ***x***(*t*), and time-independent parameters ***p*** minimizing an objective functional:

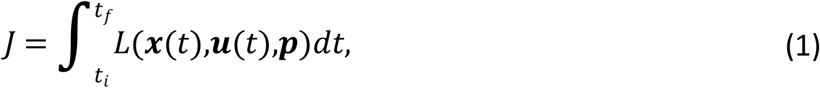

where *t_i_* and *t_f_* are initial and final times, and *t* is time. This objective functional is subject to dynamic constraints:

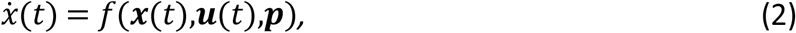

and to algebraic path constraints:

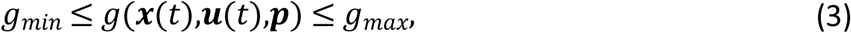

which are equality constraints if *g_min_* = *g_max_*. The optimization variables are typically bounded as follows:

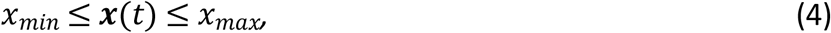

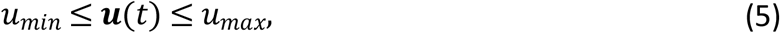

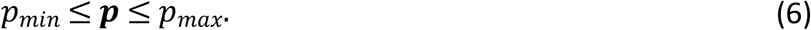

**Table 1.** Formulation of the optimal control problems.

|  | **Example 1**<br>Pendulum simulations | **Example 2**<br>2D predictive simulations of walking | **Example 3**<br>3D tracking simulations of walking |
| --- | --- | --- | --- |
| States<br>$\mathbf{x}(t)$ | Joint positions $q$ and velocities $v$ | Muscle activations $a$ and tendon forces $F_t$<br>Joint positions $q$ and velocities $v$<br>Trunk activations $a_{trunk}$ | Muscle activations $a$ and tendon forces $F_t$<br>Joint positions $q$ and velocities $v$<br>Arm activations $a_{arms}$ |
| Controls $\mathbf{u}(t)$ | Derivatives of $v$ (accelerations): $u_{dv}$<br>Joint torques $u_T$ | Derivatives of $a$: $u_{da}$<br>Derivatives of $F_t$: $u_{dF_t}$<br>Derivatives of $v$ (accelerations): $u_{dv}$<br>Trunk excitations $e_{trunk}$ | Derivatives of $a$: $u_{da}$<br>Derivatives of $F_t$: $u_{dF_t}$<br>Derivatives of $v$ (accelerations): $u_{dv}$<br>Arm excitations $e_{arms}$ |
| Parameters<br>$\mathbf{p}$ | / | Half gait cycle duration $t_f$ | Contact sphere transversal plane locations $p_{cl}$<br>Contact sphere radii $p_{cr}$ |
| Bounds | $-3\pi = q_{lb} \leq q \leq$<br>$-20 = v_{lb} \leq v \leq$<br>$-500 = u_{dv,lb} \leq$<br>$-1000 = u_{T,lb} \leq$ | $0 \leq a \leq 1$<br>$0 \leq F_t \leq F_{t,ub} = 5$<br>$-\tau_d/100 \leq u_{da} \leq \tau_a/100$<br>$\tau_d = 60\text{ms}; \tau_a = 15\text{ms}$<br>$-1 \leq u_{dF_t}, a_{trunk}, e_{trunk} \leq 1$<br>$q_{lb,man} = q_{lb} \leq q \leq q_{ub} = q_{ub,man}$<br>$v_{lb,man} = v_{lb} \leq v \leq v_{ub} = v_{ub,man}$<br>$u_{dv,lb,man} = u_{dv,lb} \leq u_{dv} \leq u_{dv,ub} = u_{dv,ub,man}$<br>$0.1 \leq t_f \leq 1$ | $0 \leq a \leq 1$<br>$0 \leq F_t \leq F_{t,ub} = 5$<br>$-\tau_d/100 \leq u_{da} \leq \tau_a/100$<br>$\tau_d = 60\text{ms}; \tau_a = 15\text{ms}$<br>$-1 \leq u_{dF_t}, a_{arms}, e_{arms} \leq 1$<br>$\hat{q}_{min} - \hat{q}_r = q_{lb} \leq q \leq q_{ub} = \hat{q}_{max} + \hat{q}_r$<br>$\hat{v}_{min} - \hat{v}_r = v_{lb} \leq v \leq v_{ub} = \hat{v}_{max} + \hat{v}_r$<br>$\hat{u}_{dv,min} - \hat{u}_{dv,r} = u_{dv,lb} \leq u_{dv} \leq u_{dv,ub} = \hat{u}_{dv,max} + \hat{u}_{dv,r}$<br>$\hat{p}_{cl} - 0.025 = p_{cl,lb} \leq p_{cl} \leq p_{cl,ub} = \hat{p}_{cl} + 0.025$<br>$\hat{p}_{cr} - 0.5\hat{p}_{cr} = p_{cr,lb} \leq p_{cr} \leq p_{cr,ub} = \hat{p}_{cr} + 0.5\hat{p}_{cr}$ |
| Scaling | $s_q = 3; \tilde{q} = q/s_q$<br>$\tilde{q}_{lb} = q_{lb}/s_q$<br>$\tilde{q}_{ub} = q_{ub}/s_q$<br>$s_t = 0.2; \tilde{v} = v/(s_q/s_t)$<br>$\tilde{v}_{lb} = v_{lb}/(s_q/s_t)$<br>$\tilde{v}_{ub} = v_{ub}/(s_q/s_t)$<br>$\tilde{u}_{dv} = u_{dv}/(s_q/s_t^2)$<br>$\tilde{u}_{dv,lb} = u_{dv,lb}/(s_q/s_t^2)$<br>$u_{dv,ub} = u_{dv,ub}/(s_q/s_t^2)$<br>$s_T = 500; \tilde{u}_T = u_T/s_T$<br>$\tilde{u}_{T,lb} = u_{T,lb}/s_T$<br>$\tilde{u}_{T,ub} = u_{T,ub}/s_T$ | $s_q = \max(\text{abs}(q_{lb}), \text{abs}(q_{ub}))$<br>$\tilde{q} = q/s_q; \tilde{q}_{lb} = q_{lb}/s_q; \tilde{q}_{ub} = q_{ub}/s_q$<br>$s_v = \max(\text{abs}(v_{lb}), \text{abs}(v_{ub}))$<br>$\tilde{v} = v/s_v; \tilde{v}_{lb} = v_{lb}/s_v; \tilde{v}_{ub} = v_{ub}/s_v$<br>$s_{u_{dv}} = \max(\text{abs}(u_{dv,lb}), \text{abs}(u_{dv,ub}))$<br>$\tilde{u}_{dv} = u_{dv}/s_{u_{dv}}$<br>$\tilde{u}_{dv,lb} = u_{dv,lb}/s_{u_{dv}}; \tilde{u}_{dv,ub} = u_{dv,ub}/s_{u_{dv}}$<br>$s_{da} = s_{dF_t} = 100; s_{trunk} = 150$<br>$s_{F_t} = F_{t,ub}; \tilde{F}_t = F_t/s_{F_t}; \tilde{F}_{t,ub} = F_{t,ub}/s_{F_t}$ | $s_q = \max(\text{abs}(q_{lb}), \text{abs}(q_{ub})); \tilde{q} = q/s_q$<br>$\tilde{q}_{lb} = q_{lb}/s_q; \tilde{q}_{ub} = q_{ub}/s_q$<br>$s_v = \max(\text{abs}(v_{lb}), \text{abs}(v_{ub})); \tilde{v} = v/s_v$<br>$\tilde{v}_{lb} = v_{lb}/s_v; \tilde{v}_{ub} = v_{ub}/s_v$<br>$s_{u_{dv}} = \max(\text{abs}(u_{dv,lb}), \text{abs}(u_{dv,ub})); \tilde{u}_{dv} = u_{dv}/s_{u_{dv}}$<br>$\tilde{u}_{dv,lb} = u_{dv,lb}/s_{u_{dv}}; \tilde{u}_{dv,ub} = u_{dv,ub}/s_{u_{dv}}$<br>$s_{da} = s_{dF_t} = 100; s_T = s_{arms} = 150$<br>$s_{F_t} = F_{t,ub}; \tilde{F}_t = F_t/s_{F_t}; \tilde{F}_{t,ub} = F_{t,ub}/s_{F_t}$<br>$\tilde{p}_{cl,v} = 1/(p_{cl,ub} - p_{cl,lb})$<br>$\tilde{p}_{cl,r} = 0.5 - p_{cl,ub}/(p_{cl,ub} - p_{cl,lb})$<br>$\tilde{p}_{cr,v} = 1/(p_{cr,ub} - p_{cr,lb})$<br>$\tilde{p}_{cr,r} = 0.5 - p_{cr,ub}/(p_{cr,ub} - p_{cr,lb})$<br>$\tilde{p}_{cl,lb} = \tilde{p}_{cr,lb} = -0.5; \tilde{p}_{cl,ub} = \tilde{p}_{cr,ub} = 0.5$ |
| Objective function | $L = w_1 \ \tilde{u}_T\ _2^2 + L_p$<br>$L_p = w_2 \ \tilde{u}_{dv}\ _2^2$ | $L = (w_1 \ a\ _3^3 + w_2 \ e_{trunk}\ _2^2 + w_3 \ \tilde{u}_{dv}\ _2^2)$<br>$L_p = 0.001 (\ u_{da}\ _2^2 + w_4 \ u_{dF_t}\ _2^2)$ | $L = w_1 \ a\ _2^2 + w_2 \ q - \hat{q}\ _2^2 + w_3 \ GRF - \hat{GRF}\ _2^2 + w_4 \ GR$<br>$L_p = 0.001 (\ u_{da}\ _2^2 + \ u_{dF_t}\ _2^2 + \ u_{dv}\ _2^2)$ |
| Dynamic constraints | $(dq/dt)/s_q = v/s_t$<br>$(dv/dt)/(s_q/s_t) = u_{dv}/(s_q/s_t^2)$ | $da/dt = s_{da} u_{da}$<br>$(dF_t/dt)/s_{F_t} = (s_{dF_t} u_{dF_t})/s_{F_t}$<br>$(dq/dt)/s_q = v/s_q$<br>$(dv/dt)/s_v = u_{dv}/s_v$ | $da/dt = s_{da} u_{da}$<br>$(dF_t/dt)/s_{F_t} = (s_{dF_t} u_{dF_t})/s_{F_t}$<br>$(dq/dt)/s_q = v/s_q$<br>$(dv/dt)/s_v = u_{dv}/s_v$ |

| | | $da_{\text{trunk}}/dt = (e_{\text{trunk}} - a_{\text{trunk}})/\tau; \tau = 35\text{ms}$ | $da_{\text{arms}}/dt = (e_{\text{arms}} - a_{\text{arms}})/\tau; \tau = 35\text{ms}$ |
| --- | --- | --- | --- |
| Path constraints | $T = f_s(q, v, u_{dv})$<br>$T/s_T = \tilde{u}_T$<br>$\tilde{q}(0) = \tilde{q}(1) = \tilde{v}(0) = \tilde{v}(1)$ | $0 \leq s_{da}u_{da} + a/\tau_d$<br>$s_{da}u_{da} + a/\tau_a \leq 1/\tau_a$<br>$f_c(a, F_t, u_{dF_t}) = 0$<br>$T = f_s(q, v, u_{dv})$<br>$T_{\text{pelvis}} = 0$<br>$T_{\text{ll}} = \sum_{m=1}^M MA_m F_{t,m}$<br>$T_{\text{trunk}}/s_{\text{trunk}} = a_{\text{trunk}}$<br>$\bar{x}(t_f) = \bar{x}(0)$<br>$(q_{\text{pelvis, for}}(t_f) - q_{\text{pelvis, for}}(0))/t_f = 1.33$ | $0 \leq s_{da}u_{da} + a/\tau_d$<br>$s_{da}u_{da} + a/\tau_a \leq 1/\tau_a$<br>$f_c(a, F_t, u_{dF_t}) = 0$<br>$T = f_s(q, v, u_{dv})$<br>$T_{\text{pelvis}}/s_T = \hat{T}_{\text{pelvis}}/s_T$<br>$T_{\text{ll, trunk}} = \sum_{m=1}^M MA_m F_{t,m} + T_p$<br>$T_{\text{arms}}/s_{\text{arms}} = a_{\text{arms}}$ |

In the first example, we perturbed the balance of 9 inverted pendulums, with between 2 and 10 degrees of freedom (DOFs), by applying a backward translation to their base of support. The optimal control problem identified the joint torques necessary to restore the pendulums’ upright posture within one second while minimizing the actuator effort (i.e., squared joint torques) and satisfying the pendulum dynamics.

In the second example, we performed predictive simulations of walking with a two-dimensional (2D) musculoskeletal model (10 DOFs, 18 muscles actuating the lower limbs, one ideal torque actuator at the trunk, and 2 contact spheres per foot [15]). We identified muscle excitations and walking cycle duration that minimized a weighted sum of muscle fatigue (i.e., muscle activations at the third power [5]) and joint accelerations subject to constraints describing the musculoskeletal dynamics, imposing left-right symmetry, and prescribing gait speed (i.e., distance travelled by the pelvis divided by gait cycle duration).

In the third example, we performed tracking simulations of walking with a three-dimensional (3D) musculoskeletal model (29 DOFs, 92 muscles actuating the lower limbs and trunk, 8 ideal torque actuators at the arms, and 6 contact spheres per foot [3,15,23]) while calibrating the foot-ground contact model. We identified muscle excitations and contact sphere parameters (locations and radii) that minimized a weighted sum of muscle effort (i.e., squared muscle activations) and the difference between measured and simulated variables (joint angles and torques, and ground reaction forces and torques) while satisfying the musculoskeletal dynamics. Data collection was approved by the Ethics Committee at UZ / KU Leuven (Belgium).

In these examples, we modeled pendulum/skeletal movement with Newtonian rigid body dynamics and, for the walking simulations, compliant Hunt-Crossley foot-ground contact [15,17]. We created a continuous approximation of a contact model from Simbody to provide twice continuously differentiable contact forces, which are required when using second-order gradient-based optimization algorithms [24]. We performed the approximations of conditional *if-tests* using hyperbolic tangent functions. For the muscle-driven walking simulations, we described muscle activation and contraction dynamics using Raasch’s model [8,25] and a Hill-type muscle model [9,26], respectively. We defined muscle-tendon lengths, velocities, and moment arms as a function of joint positions and velocities using polynomial functions [27]. We optimized the polynomial coefficients to fit muscle-tendon lengths and moment arms (maximal root mean square deviation: 3 mm; maximal order: ninth) obtained from OpenSim for a wide range of joint positions and velocities.

We transcribed each optimal control problem into a nonlinear programming problem (NLP) using a third order Radau quadrature collocation scheme. We formulated each problem in MATLAB using CasADi and IPOPT. We imposed an NLP relative error tolerance of 1 × 10^−6^ and used an adaptive barrier parameter update strategy. We selected a number of mesh intervals for each problem such that the results were qualitatively similar when using a mesh twice as fine. We used 10 and 3 initial guesses for the pendulum and walking simulations, respectively. We ran all simulations on a single core of a standard laptop computer with a 2.9 GHz Intel Core i7 processor.

### Results analysis

We compared CPU time and number of iterations required to solve the problems using the different computational choices. First, we compared AD, using the Recorder approach, with FD. Second, we compared the AD approaches, namely AD-Recorder and AD-ADOLC. We performed these two comparisons using the linear solver mumps [28], which CasADi provides, and an approximated Hessian. Third, we compared different linear solvers, namely mumps with the collection of solvers from HSL [29], while using AD-Recorder and an approximated Hessian. Finally, we compared the use of approximated and exact Hessians. For this last comparison, we used AD-Recorder and tested all linear solvers. In all cases, we ran simulations from different initial guesses and compared results from simulations that started from the same initial guess and converged to similar optimal solutions.

## Results

Using AD-Recorder was computationally more efficient than using FD or AD-ADOLC (Fig 3). The CPU time decreased when using AD-Recorder as compared with FD (between 1.8 ± 0.1 and 17.8 ± 4.9 times faster with AD-Recorder) and AD-ADOLC (between 3.6 ± 0.3 and 12.3 ± 1.3 times faster with AD-Recorder). CPU time spent in evaluating the objective function gradient accounted for 95 ± 10% (average ± standard deviation) of the difference in CPU time between AD-Recorder and FD. The difference in CPU time spent in evaluating the constraint Jacobian accounted for 89 ± 6% of the difference in CPU time between AD-Recorder and AD-ADOLC. The number of iterations was similar when using AD-Recorder, FD, and AD-ADOLC. For the 2D predictive and 3D tracking simulations, 1 and 2 cases, respectively, out of 9 (3 derivative scenarios and 3 initial guesses) were excluded from the comparison as they converged to different solutions.

**Fig 3.**
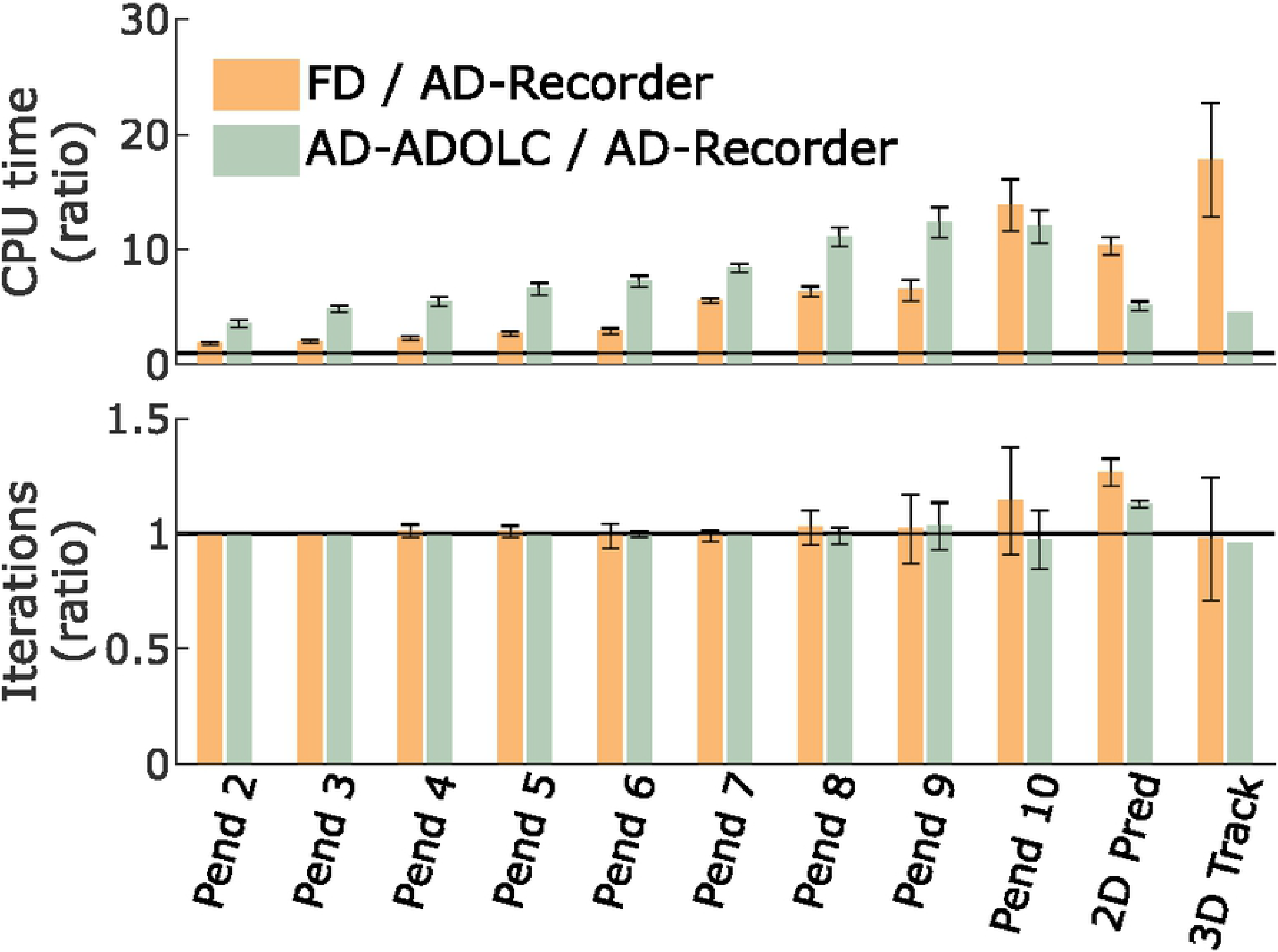
Comparison of CPU time (top) and number of iterations (bottom) between FD, AD-ADOLC, and AD-Recorder. The comparisons are expressed as ratios and averaged over results from different initial guesses (error bars represent ± one standard deviation). The horizontal lines indicate 1:1 ratios. Ratios larger than one indicate slower convergence (top) and more iterations (bottom) with FD or AD-ADOLC as compared to AD-Recorder. Pend indicates pendulum simulations with the number being the number of DOFs; Pred and Track indicate predictive and tracking simulations, respectively. The results were obtained using mumps and an approximated Hessian.

The solvers from the HSL collection were on average more efficient (faster with a similar number of iterations) than mumps for the pendulum simulations, but the efficiency varied for the 2D predictive and 3D tracking simulations (Table 2). The solver ma27 was on average faster than mumps in all cases although ma27 required more iterations for the 2D predictive simulations. The other solvers from the HSL collection were on average slower than mumps for the 2D predictive simulations. For the 3D tracking simulations, the solvers ma77 and ma86 were faster and slower, respectively, than mumps. The solvers ma57 and ma97 failed to solve the 3D tracking simulations due to memory issues. For all simulations, the solvers from the HSL collection except ma86 (and ma77 for the 2D predictive simulations) required less CPU time per iteration than mumps. For the 2D predictive and 3D tracking simulations, 1 case out of 18 (6 solvers and 3 initial guesses) and 4 cases out of 12 (4 solvers and 3 initial guesses), respectively, were excluded from the comparison as they converged to different solutions.

**Table 2.** Comparison of CPU time, number of iterations, and CPU time per iteration between linear solvers.

| Solver<br>* vs<br>mumps | Pendulum-based simulations |  |  | 2D predictive simulations |  |  | 3D tracking simulations |  |  |
| --- | --- | --- | --- | --- | --- | --- | --- | --- | --- |
|  | CPU time | Iteration<br>Number | CPU time<br>per<br>Iteration | CPU time | Iteration<br>Number | CPU time<br>per<br>Iteration | CPU time | Iteration<br>Number | CPU time<br>per<br>Iteration |
| *ma27 | $0.6 \pm 0.1$ | $1.0 \pm 0.0$ | $0.6 \pm 0.1$ | $1.0 \pm 0.3$ | $1.4 \pm 0.3$ | $0.7 \pm 0.1$ | $0.6 \pm 0.3$ | $0.8 \pm 0.3$ | $0.7 \pm 0.1$ |
| *ma57 | $0.6 \pm 0.0$ | $1.0 \pm 0.0$ | $0.6 \pm 0.0$ | $2.4 \pm 2.5$ | $2.8 \pm 2.8$ | $0.8 \pm 0.0$ | / | / | / |
| *ma77 | $0.9 \pm 0.1$ | $1.0 \pm 0.0$ | $0.9 \pm 0.0$ | $1.3 \pm 0.0$ | $1.2 \pm 0.0$ | $1.1 \pm 0.0$ | $0.5 \pm 0.1$ | $0.6 \pm 0.2$ | $0.8 \pm 0.0$ |
| *ma86 | $1.1 \pm 0.1$ | $1.0 \pm 0.0$ | $1.1 \pm 0.1$ | $2.1 \pm 0.5$ | $1.1 \pm 0.1$ | $1.8 \pm 0.3$ | $2.3 \pm 0.0$ | $1.7 \pm 0.0$ | $1.3 \pm 0.0$ |
| *ma97 | $0.7 \pm 0.0$ | $1.0 \pm 0.0$ | $0.7 \pm 0.0$ | $1.2 \pm 0.3$ | $1.2 \pm 0.1$ | $1.0 \pm 0.2$ | / | / | / |

The comparisons are expressed as ratios (mean ± one standard deviation; results obtained with solver from the HSL collection over results obtained with mumps). The ratios are averaged over results from different initial guesses. Ratios larger than one indicate faster convergence, fewer iterations, or more time per iteration with mumps. The use of the solvers ma57 and ma97 led to memory issues for the 3D tracking simulations and these cases were therefore excluded from the analysis. The simulations were run using AD-Recorder and an approximated Hessian.

Using an exact Hessian was more efficient than using an approximated Hessian for the pendulum simulations but not for the 2D predictive simulations (Fig 4). The exact Hessian required less CPU time and fewer iterations than the approximated Hessian for the pendulum simulations (average 2.4 ± 1.2 times faster and 2.5 ± 0.9 times fewer iterations). By contrast, the exact Hessian required more CPU time and iterations than the approximated Hessian for the 2D predictive simulations (average 6.0 ± 0.8 times slower and 2.1 ± 0.2 times more iterations). For the pendulum simulations, 27 cases out of 540 (9 pendulums, 6 solvers, and 10 initial guesses) were excluded from the comparison as they converged to different solutions with the two Hessian settings. One case was also excluded as it had not converged after 3000 iterations with the exact Hessian but converged in 209 iterations with the approximated Hessian. For the 2D predictive simulations, only results obtained with the solvers ma86 and ma97 were included, since the use of the other solvers led to memory issues. Further, 4 cases out of 6 (2 solvers and 3 initial guesses) were excluded from the comparison as they converged to different solutions with the two Hessian settings. Finally, the 3D tracking simulations were not included for this comparison as the large problem size induced memory issues with the exact Hessian.

**Fig 4.**
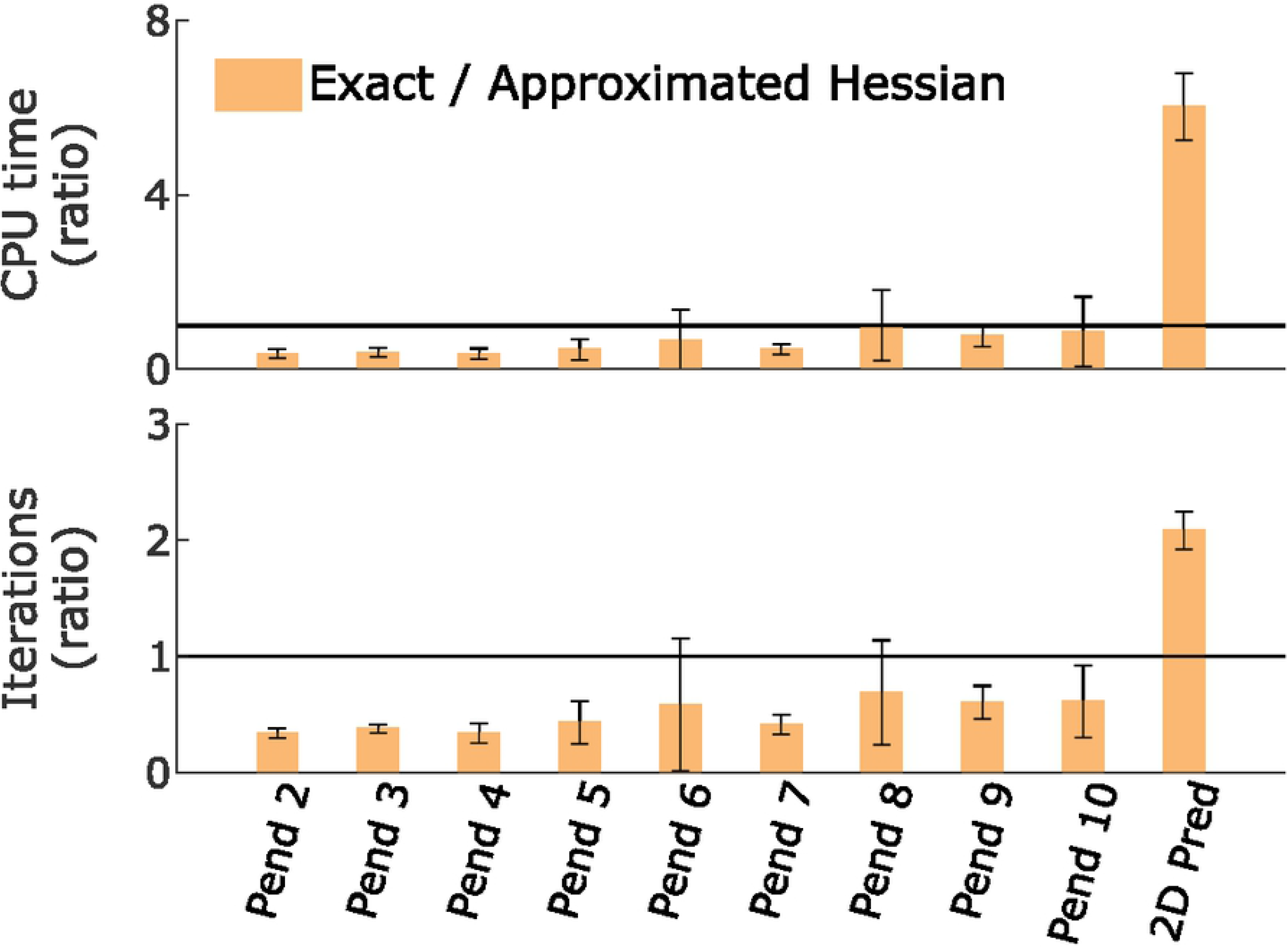
Comparison of CPU time (top) and number of iterations (bottom) between exact an approximated Hessian. The comparisons are expressed as ratios and averaged over results from different initial guesses (error bars represent ± one standard deviation). The horizontal lines indicate 1:1 ratios. Ratios larger than one indicate slower convergence (top) and more iterations (bottom) with an exact versus an approximated Hessian. Pend indicates pendulum simulations with the number being the number of DOFs; Pred and Track indicate predictive and tracking simulations, respectively. The results were obtained using all solvers and AD-Recorder.

The pendulum simulations required at most 21 s and 366 iterations to converge (results obtained with AD-Recorder, mumps, and an approximated Hessian); CPU time and number of iterations depended on the number of DOFs (S1 Movie).

The 2D predictive simulations reproduced salient features of human gait but deviated from experimental data in three noticeable ways (Fig 5; S2 Movie). First, the predicted knee flexion during mid-stance was limited, resulting in small knee torques. Second, the simulations produced less ankle plantarflexion at push-off. Third, the vertical ground reaction forces exhibited a large peak at impact. The simulations converged in less than one CPU minute (average over solutions starting from three initial guesses: 36 ± 17 s and 247 ± 143 iterations; results obtained with AD-Recorder, mumps, and an approximated Hessian).

**Fig 5.**
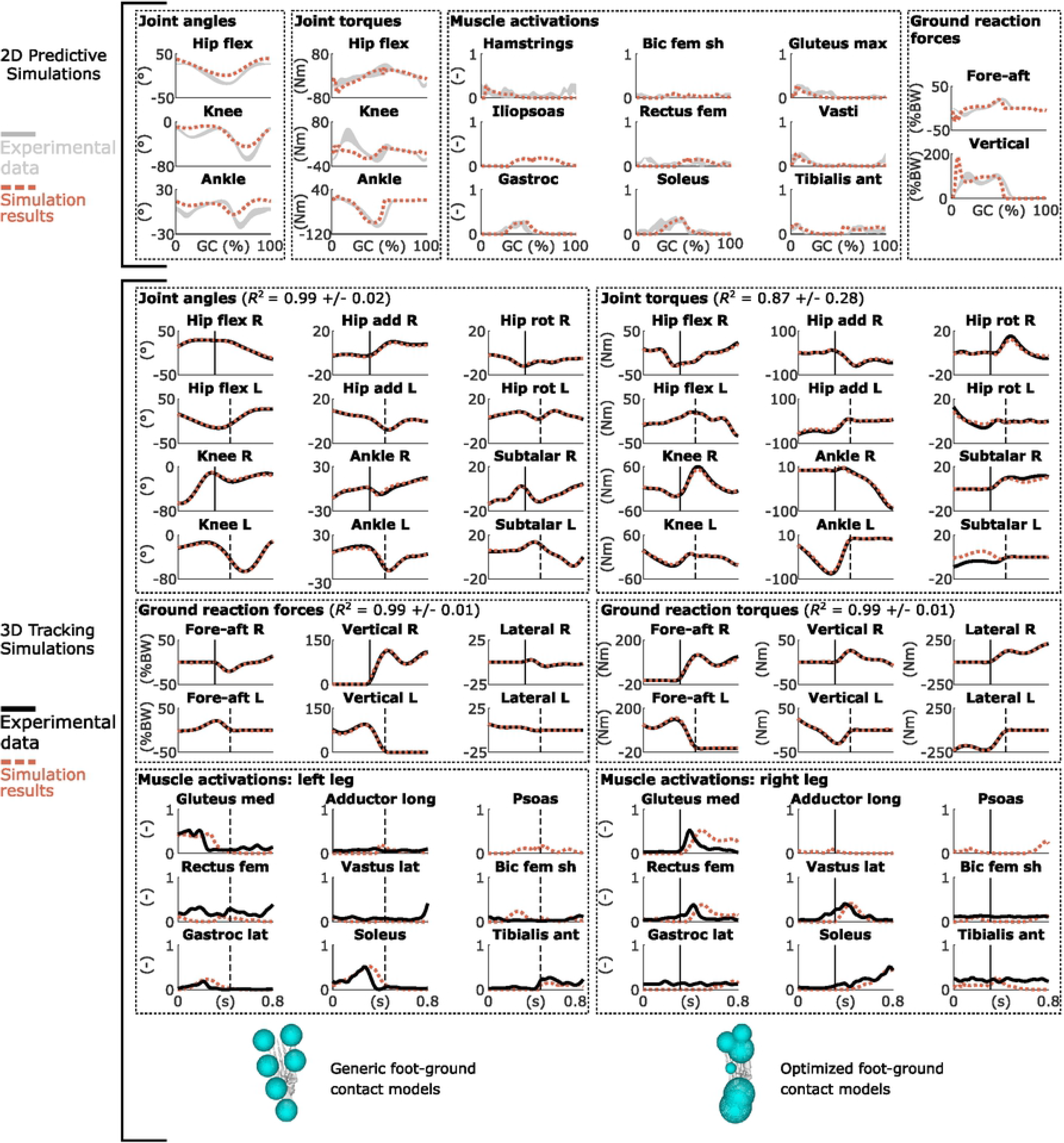
Results from optimal control simulations of walking. (Top) Results from 2D predictive simulations of walking (joint angles: flex is flexion, GC is gait cycle; muscle activations: bic is biceps, fem is femoris, sh is short head, max is maximus, gastroc is gastrocnemius, ant is anterior; ground reaction forces: BW is body weight). Experimental data are shown as mean ± two standard deviations. (Bottom) Results from 3D tracking simulations of walking (joint angles: R is right, L is left, add is adduction, rot is rotation; muscle activations: med is medialis, long is longus, lat is lateralis). The vertical lines indicate right heel strike (solid) and left toe-off (dashed); only part of the gait cycle, when experimental ground reaction forces are available, is tracked. The experimental EMG data are normalized to peak muscle activations. The foot diagrams depict a down-up view of the configuration of the contact spheres of the right foot pre-calibration (left; generic) and post-calibration (right; optimized). The coefficient of determination *R^2^* is given for the tracked variables.

The 3D tracking simulations accurately tracked the experimental walking data (average coefficient of determination *R^2^*: 0.95 ± 0.17; Fig 5; S3 Movie). Simulated muscle activations also qualitatively resembled experimental electromyography (EMG) data, even though EMG was not tracked (Fig 5). The configuration of the contact spheres differed from the generic model after the calibration. The simulations converged in less than 20 CPU minutes (average over simulations starting from two initial guesses: 19 ± 7 minutes and 493 ± 151 iterations; results obtained with AD-Recorder, mumps, and an approximated Hessian).

## Discussion

We showed that the use of AD over FD improved the computational efficiency of OpenSim-based optimal control simulations. Specifically, AD drastically decreased the CPU time spent in evaluating the objective function gradient. This time decrease results from AD’s ability to evaluate a Jacobian-transposed-times-vector product through its reverse mode. The objective function gradient has many inputs (all optimization variables) but only one output. It can thus be evaluated in only one reverse sensitivity sweep; the computational cost is hence proportional to the cost of evaluating the objective function. By contrast, with FD, the computational cost is proportional to the number of optimization variables times the cost of evaluating the objective function. The efficiency benefit of AD also increased with the complexity of the problems. This is expected, since the number of optimization variables increases with the problem size; FD thus requires more objective function evaluations, whereas AD still only requires one reverse sweep.

The choice of the objective function influences CPU time. As an illustration, we added a term representing the metabolic energy rate [30] to the objective function of the 2D predictive simulations. Minimizing metabolic energy rate is common in predictive studies of walking [4,6,24]. Solving the resulting optimal control problem was about 60 times faster with AD-Recorder than with FD (although FD required fewer iterations), whereas AD-Recorder was only about 10 times faster than FD without incorporating the metabolic energy rate in the objective function. This increased time difference can be explained by our use of computationally expensive hyperbolic tangent functions to make the metabolic energy rate model twice continuously differentiable, as required when using second-order gradient-based optimization algorithms [24]. Overall, AD reduces the number of function evaluations, which has an even larger effect if these functions are expensive to compute.

The implementation of AD was computationally more efficient through Recorder than through ADOL-C. Specifically, Recorder decreased the CPU time by a factor 4-12 compared to ADOL-C. ADOL-C records all calculations involving differential variables on a sequential data set called a tape [20], which is then evaluated by ADOL-C ’s *virtual machine*. By contrast, Recorder generates plain C-code. The factor 4-12 is the difference between a *virtual machine* interpreting a list of instructions (ADOL-C) and machine code performing these instructions directly (Recorder). In future work, we aim to support computer programs containing branches with Recorder.

It is difficult to provide guidelines for the linear solver selection based on our results, as their efficiency was problem-dependent. In contrast with mumps, the solvers from the HSL collection do not freely come with CasADi and are only free to academics. Our study hence does not support the extra effort to obtain them since they did not consistently outperform mumps in our applications. Yet an in-depth analysis of the solvers’ options and underlying mathematical details should be considered in future work.

The use of an exact Hessian, rather than an approximated Hessian, improved the computational efficiency for the pendulum simulations but not for the walking simulations. For the 2D walking simulations, using an exact Hessian required more CPU time but also more iterations. This might seem surprising, since an exact Hessian is expected to provide more accurate information and, therefore, lead to convergence in fewer iterations. However, IPOPT requires the Hessian to be positive definite when calculating a Newton step to guarantee that the step is in the descent direction. When this is not the case, the Hessian is approximated with a positive definite Hessian by adding the identity matrix multiplied by a regularization term to the Hessian [21]. We observed that for the 2D predictive simulations, the magnitude of the regularization term was much greater than for the pendulum simulations. Yet excessive regularization might degrade the performance of the algorithm, as regularization alters the second-order derivative information and causes IPOPT to behave more like a steepest-descent algorithm [31]. The approximated Hessian requires no regularization, which likely explains the difference in number of iterations. Overall, convexification of the currently non-convex optimal control problems is expected to further improve the computational efficiency [8].

The 2D predictive and 3D tracking simulations produced realistic movements although deviations remain between simulated and measured data. Modeling choices rather than local optima likely explain these deviations. These choices have a greater influence on the predictive simulations, since deviations from measured data are minimized in tracking simulations whereas only the motor task goal is specified in the objective function of predictive simulations. Several modeling choices might explain the main deviations for the predictive simulations. First, we did not model stability requirements, which might explain the limited knee flexion during mid-stance [5,24]. Instead, we included muscle activity in the cost function, which might explain why reducing knee torques and, therefore, knee extensor activity was optimal. Second, the model did not include a metatarsophalangeal (MTP) joint, which might explain the limited ankle plantarflexion at push-off; similar ankle kinematics have indeed been observed experimentally when limiting the range of motion of the MTP [32]. Third, the lack of knee flexion combined with the simple trunk model (i.e., one DOF controlled by one ideal torque actuator) might explain the high vertical ground reaction forces at impact [5]. Finally, the goal of the motor task (i.e., minimizing muscle fatigue) likely does not fully explain the control strategies governing human walking. In this study, the focus was on evaluating different computational choices but future work should exploit the improved computational efficiency to explore how modeling choices affect the correspondence between simulated and measured quantities.

## Conclusions

In this study, we enabled the use of AD when performing OpenSim-based optimal control simulations. We showed that using AD drastically improved the computationally efficiency of such simulations. This improved efficiency is highly desirable for researchers using complex models or aiming to implement such models in clinical practice where time constraints are typically more stringent than in research context. Overall, the combination of AD with other efficient numerical tools such as direct collocation and implicit differential equations allows overcoming the computational roadblocks that have long limited the use of optimal control simulations for biomechanical applications. In the future, we aim to exploit this computational efficiency to design optimal treatments for neuro-musculoskeletal disorders, such as cerebral palsy.

## Acknowledgments

The authors would like to thank Michael Sherman for helpful technical discussions.

## Supporting information

**S1 Appendix. Example source code.** Recorder provides the expression graph of the function to differentiate as MATLAB source code in a format that CasADi’s AD algorithms can then transform into C-code. This file provides MATLAB and C source code resulting from applying these two steps on the example function from Fig 1.

**S1 Movie. Pendulum-based simulations of perturbed balance**. The pendulums have between 2 and 10 degrees of freedom. The playback speed is 0.2 times real-time.

**S2 Movie. 2D muscle-driven predictive simulation of walking (1.33 m s^−1^)**. The playback speed is 0.2 times real-time.

**S3 Movie. 3D muscle-driven tracking simulation of walking**. The pink model (tracking simulation results) tracks the motion of the white model (experimental data). The playback speed is 0.2 times real-time.

